# The serotonin 1B receptor is required for some of the behavioral effects of psilocybin in mice

**DOI:** 10.1101/2024.10.18.618582

**Authors:** Sixtine Fleury, Katherine M. Nautiyal

**Author notes:** To whom correspondence should be addressed 6307 Moore Hall Hanover, NH 03755.

## Abstract

Recent studies highlight the promising use of psychedelic therapies for psychiatric disorders, including depression. The persisting clinical effects of psychedelics such as psilocybin are commonly attributed to activation of the serotonin 2A receptor (5-HT2AR) based on its role in the acute hallucinatory effects. However, the active metabolite of psilocybin binds to many serotonin receptor subtypes, including the serotonin 1B receptor (5-HT1BR). Given the known role of 5-HT1BR in mediating depressive phenotypes and promoting neural plasticity, we hypothesized that it mediates the effects of psilocybin on neural activity and behavior. We first examined the acute neural response to psilocybin in mice lacking 5-HT1BR. We found that 5-HT1BR expression influenced brain-wide activity following psilocybin administration, measured by differences in the patterns of the immediate early gene c-Fos, across regions involved in emotional processing and cognitive function, including the amygdala and prefrontal cortex. Functionally, we demonstrated that 5-HT1BR mediates some of the acute and persisting behavioral effects of psilocybin. Although there was no effect of 5-HT1BR expression on the acute head twitch response, mice lacking 5-HT1BRs had attenuated hypolocomotion to psilocybin. We also measured the persisting effects of psilocybin on anhedonia and anxiety-like behavior using transgenic and pharmacological 5-HT1BR loss-of-function models and found that 5-HT1B is involved in mediating the decreased anhedonia and reduced anxiety-like behavior. Finally, using a network analysis, we identified neural circuits through which 5-H1BR may modulate the response to psilocybin. Overall, our research implicates the 5-HT1BR, a non-hallucinogenic serotonin receptor, as a mediator of the behavioral and neural effects of psilocybin in mice.

## Introduction

In recent years, almost 10% of adults in the US meet the criteria for a mood disorder, and nearly 20% for an anxiety disorder (1). Despite the widespread use of selective serotonin reuptake inhibitors (SSRIs) as antidepressants and anxiolytics, these treatments require long-term administration and fail to provide relief for a significant proportion of patients (2). Developing novel therapeutic targets which include rapid-acting effective therapies is important for the treatment of depressive and anxiety disorders. Psychedelic compounds are emerging as potential therapeutic agents for treating psychiatric disorders. Growing evidence supports their efficacy in treating major depressive disorder (MDD) and other mood and anxiety disorders (3–8). Psilocybin is a serotonergic psychedelic found in *Psilocybe* mushrooms which is gaining traction as a promising therapeutic agent (9). Clinical studies report that one to two doses of psilocybin can produce antidepressant effects in MDD and treatment resistant depression patients that persist for weeks to months (5,7,10–14). However, the hallucinogenic properties of psilocybin limit its use for some patient populations and in everyday clinical practice. Uncovering the neural mechanisms responsible for the therapeutic benefits in preclinical studies is therefore crucial to developing other treatment strategies (15,16).

The persisting behavioral effects have been largely attributed to the activation of the 5-HT2A receptor, based on its well-documented role in mediating the acute hallucinogenic properties of serotonergic psychedelics (17–21). However, there is conflicting evidence emerging from preclinical studies on the necessity of 5-HT2AR in the behavioral effects, especially related to depressive behavior using pharmacological and genetic loss-of-function models (22–28). Some recent studies suggest that these behavioral and neural effects may be independent of 5-HT2AR activation (23,27). Psilocybin’s capacity to drive neuroplastic changes has been demonstrated through structural changes in dendritic spines, including increased spine density and size in the prefrontal cortex and synaptic changes in the hippocampus (29) —all alterations which may remain intact with blockade of psilocybin binding to 5-HT2A receptors (23,30). Recent studies also implicate the neurotrophin receptor TrkB and brain-derived neurotrophic factor (BDNF) signaling in mediating psilocybin-induced synaptic plasticity, independent of 5-HT2ARs (31).

Psilocybin’s active metabolite, psilocin, binds with high affinity to many serotonin receptor subtypes, not just 5-HT2A, and its polypharmacology is likely important for its clinical efficacy (32). One receptor besides 5-HT2A that binds psilocybin and may be important for the effects of psilocybin on behavior is the serotonin 1B receptor (5-HT1BR). As an inhibitory Gi/o-coupled G protein-coupled receptor (GPCR) that inhibits neurotransmitter release at axon terminals, it has been implicated in a number of plasticity mechanisms including long-term potentiation of cortical inputs to the hippocampus, and long-term depression of corticostriatal synapses (33,34). 5-HT1BRs have also been linked to increased anxiety and depressive behaviors (35–37). Finally, 5-HT1BR has been implicated in mediating the effects of other psychoactive substances including MDMA and Ketamine (38–42). These data suggest that 5-HT1B could be a critical mediator of psilocybin’s persisting therapeutic effects.

Here, we hypothesize that psilocybin modulates depressive-like behaviors through 5-HT1B receptor activation. Using a genetic and pharmacological loss-of-function models, we sought to investigate the role of 5-HT1B in neural and behavioral effects of psilocybin in mice. We assessed the role of 5-HT1BR in modulating the acute drug effects of psilocybin on brain-wide neural activity, head twitch, and locomotor behavior. We also investigated the impact of 5-HT1BR on the long-term behavioral effects of psilocybin on anxiety and anhedonia phenotypes, key features of depressive disorders. Our findings suggest that the 5-HT1BR influences brain-wide neural changes following psilocybin administration and is necessary for its enduring antidepressant-like effects in mice

## Materials and Methods

### Animals

Mice were bred in the Dartmouth College vivarium and weaned at postnatal day (PN) 21 into cages of 2–4 same-sex littermates. Cohorts of mice lacking 5-HT1BR expression and littermate controls were generated by crossing female homozygous floxed-tetO1B mice to male homozygous floxed-tetO1B mice with the βActin-tTS transgene (tetO1B+/+ females crossed to tetO1B+/+::βActin-tTS + males), as previously reported (43). N=27 (11 males, 16 females) were used in the c-Fos study. N=52 mice (31 females, 21 males) were used to assess the acute behavioral effects of psilocybin. N=155 (60 males, 95 females) were used to assess persisting behavioral effects of psilocybin following chronic corticosterone administration. N=164 mice (61 females, 103 males) were used to assess persisting behavioral effects of psilocybin following chronic behavioral despair. Mice were run in 3–4 experimental cohorts, and all treatment conditions were represented within each cohort. N = 43 female mice were used for the antagonist experiment. For the adult knockout experiment (N=30), temporal-specific knockdown was achieved by withdrawal of doxycycline (Product # F5545, Bio-Serv, Flemington, NJ) at PN60 from mice following administration to the dam throughout gestation and nursing, and to the mice following weaning. As previously reported, this results in normal levels of 5-HT1BR expression through development and knockout in adulthood (43). Following PN120, mice were then administered chronic corticosterone administration and persisting behavioral effects of psilocybin were examined. All mice were maintained on a 12:12 light-dark cycle and on *ad libitum* chow and water except as noted below for treatments and behavioral testing. Mice were between ages 3-9 months old at time of testing. All procedures were approved by the Dartmouth College Institutional Animal Care and Use Committee.

### Drugs and treatments

Psilocybin (Cat # 14041, Cayman Chemical), or saline vehicle, was injected intraperitoneally (i.p.) at the dose of 5mg/kg diluted in saline solution (0.9% NaCl) at a volume of 10ml/kg. The dose of 5mg/kg was chosen based on our pilot experiments in this strain of mice that induced post-acute behavioral changes. Although the dose is on the higher side of the usual range of 1-5mg/kg commonly used in rodents, is in line with other reports assessing behavioral effects of psilocybin in mice with doses over 4mg/kg (44–46). For antagonist experiments, 0.9% saline vehicle, or the serotonin 5-HT1B/1D receptor antagonist GR127935 (CAT#508014; Sigma-Aldrich) was injected i.p. at 10mg/kg (in 0.9% saline at 10ml/kg volume) 30 minutes prior to psilocybin or saline vehicle administration, as previously used for pretreatment in behavioral pharmacology studies to target 5-HT1B/D receptors (47–50).

#### Chronic corticosterone administration

Corticosterone (35 μg/mL; Cat # 2505, Sigma-Aldrich) was dissolved in 0.45% beta-cyclodextrin (Cat # 4767, Sigma-Aldrich) in drinking water, and delivered *ad libitum* in opaque bottles for 4 weeks. This paradigm produces a corticosterone dosing of approximately 9.5 mg/kg/day (51). In non-stressed mice, only 0.45% beta-cyclodextrin was dissolved in drinking water, and delivered *ad libitum* in opaque bottles for 4 weeks.

#### Chronic behavioral despair model

As an alternative to chronic corticosterone administration, a separate group of male mice were subjected to repeated stress exposures via five forced swim sessions. Each mouse was placed in a 2-liter beaker containing water (22-25°C) for 10 minutes daily for five consecutive days. Mice remained in their home cages between sessions. Mice were injected with psilocybin (1mg/kg, i.p) or saline vehicle (10 ml/kg) 48 hours after the last swim session.

### Behavioral tests

#### Head Twitch Response and Locomotion

Head twitches and locomotion were analyzed in a subset of mice after the administration of either vehicle or psilocybin (5mg/kg, i.p) in both control and mice lacking 5-HT1BR. Immediately following injections, mice were placed in an open field, and video recording was initiated with a camera positioned directly above the arena for 1 hour. Three experimenters scored the first 15 minutes of activity for head twitches while blinded to treatment, noting the total number of and time stamp of each head twitch to compare across experimenters. Locomotion was scored with EthoVision software.

#### Gustometer

A Davis Rig 16-bottle Gustometer (Med Associates MED-DAV-160 M) was used to test the effects of psilocybin and 5-HT1BR expression on hedonic responding. Mice were water-restricted for 5 days of initial training before being put on corticosterone. On day 1 of training, mice were placed in the gustometer as a cage and with free access to 5% sucrose for 15 minutes. On day 2, mice were placed individually with free access to 5% sucrose for 15 minutes. On day 3, mice were placed individually with access to 5% and 10% sucrose with tubes alternating for 30 minutes. Mice were then food restricted for 2 days before being individually returned to the gustometer for 30 minutes with access to water, 2% sucrose, 4% sucrose, 6% sucrose, 8% sucrose, and 10% sucrose for a final training day using 6 bottle paradigm. Each concentration was randomly presented for a maximum of 120 seconds at a time before switching to another tube. Total licks at each tube were recorded using a capacitance-based system. Mice that had less than 300 licks during the final training session were excluded from the analysis. After the final training session, mice were put on corticosterone. The 6 bottle paradigm was then repeated 24 hours after drug administration for the experimental gustometer test. Each mouse was weighed before the 6 bottle paradigm and data were normalized by weight. After the experimental test, mice were allowed food that day until 24 hours before the time of the novelty-suppressed feeding test (about ∼1h30).

#### Novelty Suppressed Feeding (NSF)

The NSF assay was conducted in a rodent transport container (20’ x 16’) with the floor covered with approximately 2-4cm of corn bedding to serve as an arena. Thirty minutes before testing, mice were placed individually into holding cages. At the time of testing, a single pellet of food was placed on a white platform (∼4’ diameter) in the center of the arena. The anxiogenic environment was produced by placing a lamp with high luminosity (∼120 lux) above the center of the arena. Each mouse was placed in the same corner of the arena, facing the wall, and a stopwatch was immediately started. The latency to eat, defined as the mouse biting the pellet, was recorded over a session of 5 minutes. Immediately after a bite, the pellet was removed from the arena. To control for hunger, mice were immediately placed in their homecage after the testing session with a new food pellet, and the stopwatch was started again to measure latency to eat within their homecage. Animals that did not bite during the testing session were attributed the maximum value (300 seconds).

Animals were placed back in their homecage with food and water ad libitum once all cage mates were tested. The NSF was performed 24 hours after the gustometer in the corticosterone stress paradigm, and 48 hours after the EPM in the chronic behavioral despair paradigm.

#### Elevated Plus Maze (EPM)

Mice were placed in the center facing a closed arm and videotaped while allowed to explore the maze undisturbed for 6 minutes. The maze has four arms (35cm long x 5cm wide) that were 60cm above the ground. The two opposite closed arms had 20cm high walls, while the two opposing open arms had a 1cm lip.

Behavior was scored using EthoVision software for the number of entries and time spent in each arm. Mice were excluded from the analysis as outliers if the number of open arm entries was greater than or less than two standard deviations from the mean. The EPM was performed 24 hours after the NSF in the corticosterone stress paradigm, and 24 hours after the gustometer in the chronic behavioral despair paradigm.

### c-Fos Whole Brain Analysis

#### Brain samples

A set of male and female tTS-(N=14) and tTS+ (N=14) mice not maintained on doxycycline was used to perform a whole brain c-Fos analysis following vehicle or psilocybin (5mg/kg, i.p) administration. Brain-wide c-Fos expression was induced during the acute phase of psilocybin administration. Mice were injected with either vehicle or psilocybin (5mg/kg), and then placed back in their home cage for 2h, before perfused with 0.1 M PBS followed by 4% formaldehyde. Brains were extracted and postfixed for 24 hours in 4% formaldehyde before being cryoprotected in 30% sucrose. Brains were then sectioned in a coronal plane with a thickness of 40um on a cryostat (Leica Biosystems) and stored in 0.1 M PBS at 4°C in 24 well plates. About 20 to 30 sections per brain were selected for staining. One brain in the saline-WT group was eliminated due to technical issues with tissue sections prior to processing.

#### Tissue processing

Sections were first washed three times in PBS-T (0.1%) for 10 minutes. Sections were then blocked for 1h in 2% Normal Donkey Serum (#NC9624464, Fisher) in PBS-Triton (0.1%). Sections were then incubated in primary antibody (rb anti-cfos Rabbit Recombinant Monoclonal c-Fos antibody, ab214672) at 1:1000 in blocking solution overnight at 4°C. The following day, sections were washed 3 times again in PBS-T (0.1%). Sections were then incubated in secondary antibody Alexa Fluor 488 anti-rabbit (ab150073, Abcam) at 1:250 and DAPI at dilution 1:10,000 in PBT (0.15) for 2 hours away from light. Sections were then washed again three times and mounted.

Once dry, the slides were cover slipped with ProLong Gold. Whole sections were then imaged on Keyence BZ-X800 microscope with a 10x objective. Fluorophores were imaged with DAPI and GFP filter cubes.

#### Segmentation of fluorescent labels and quantification

Cells expressing a c-Fos label were segmented using the machine learning-based pixel and object classification program, *Ilastik*, as previously described in Berg et al., 2019. *Ilastik* was also used to generate binary images of the segmented c-Fos labels. Registration and quantification of histological images on a brain-wide scale were performed using the open-source atlas registration tool FASTMAP, as previously described in (52). Regions that were not adequately aligned or identified on stained sections were left out of the overall analysis.

## Statistical Analyses

All behavioral analyses were performed using GraphPad Prism (v.10.2.2) or RStudio (v.2023.06.1+524). The threshold of statistical significance was set at 0.05. Data were analyzed using one-, two-, and three-ANOVAs, linear mixed models, generalized linear models, and t-tests using GraphPad Prism and RStudio, with Tukey’s correction for multiple comparisons when appropriate. Area under the curve for locomotion was calculated using the trapz function in RStudio. Functional connectivity analyses were performed on a collection of 30 regions based on our interest and ability to reliably and consistently delineate these regions using a DAPI stained reference image. A psilocybin index was generated by subtracting average total c-FOS counts. The density of regional c-Fos expression was cross-correlated within each group to generate pairwise correlation matrices, and difference matrices were obtained by subtracting coefficients within each element. To quantify psilocybin-induced changes in network synchrony within each genotype, we extracted the upper triangle to obtain pairwise correlation changes. We then performed a one-sample-t-test on each vector to determine whether the mean change in correlation significantly differed from zero within each genotype. The similarity and strength of matrices were assessed using nonparametric permutation testing Monte Carlo and Spearman correlation to generate rho and p values. For network analysis, we performed a fisher r-to-z transformation on correlation and selected significant edges at 95%, 90%, 85%, and 80% confidence intervals.

## Results

### Psilocybin-induced changes in neural activity require 5-HT1BR expression

To investigate how psilocybin changes brain-wide neural activity, and whether 5-HT1BR expression influences those changes, we performed a whole-brain c-Fos analysis on male and female mice lacking the 5-HT1BR and their littermate controls. Brains were collected 2h following a single injection of psilocybin (5mg/kg) or saline vehicle, and we found that the number of c-Fos+ cells was relatively consistent across animals within brain regions (Fig 1A). Overall, there was a significant effect of psilocybin on neural activity measured by c-Fos staining (main effect of psilocybin: F(1,23)=5.708, p=0.026). Interestingly, the neural activity was significantly influenced by 5-HT1BR expression (genotype by drug interaction: F(1, 23)=5.986, p=0.023), though the pattern varied by brain region (Fig S1; genotype by drug by region interaction: F(28, 628) = 2.352, p=0.0001). Some brain regions showed significant effects of psilocybin regardless of 5-HT1BR expression, such as the somatosensory cortex and the claustrum (CLA) which both express high levels of 5-HT2ARs (Fig 1B, main effect of drug on CLA: F(1, 22)=21.74, p=0.0001). There were also select brain regions, such as the basolateral amygdala (BLA) which showed differential baseline c-Fos activity based on genotype (main effect of genotype: F(1, 22) = 4.832, p = 0.039) which potentially drove the differential effect of psilocybin between genotypes (genotype by drug interaction: F(1, 22) = 11.13, p = 0.003). Most interestingly, quite a few other brain regions showed an effect of 5-HT1BR expression on the c-Fos response to psilocybin including prefrontal cortical areas and the central amygdala (CEA) which showed increased c-Fos in control, but not 5-HT1BR KO mice (drug by genotype interaction for CEA: F(1, 23)=8.084, p=0.009; controls: p=0.0002, 5-HT1BR KOs: p=0.669). The globus pallidus showed a similar pattern with increased c-Fos expression in controls, but not 5-HT1BR KO mice (drug by genotype interaction for GP: F(1, 22)=4.954, p=0.037; controls: p=0.003; 5-HT1BR KO: p=0.829). Importantly, these regions express high levels of 5-HT1BR protein (Fig S2; (53,54), and may be key sites for psilocybin acting via these receptors to influence neural activity. Overall, psilocybin differentially influenced neural activity throughout the brain leading to both increases and decreases depending on the brain area, and these changes in neural activity were differentially influenced by 5-HT1BR expression (Fig 1C). This points to a role for the 5-HT1BR to mediate the behavioral effects of psilocybin in rodents.

**Figure 1:**
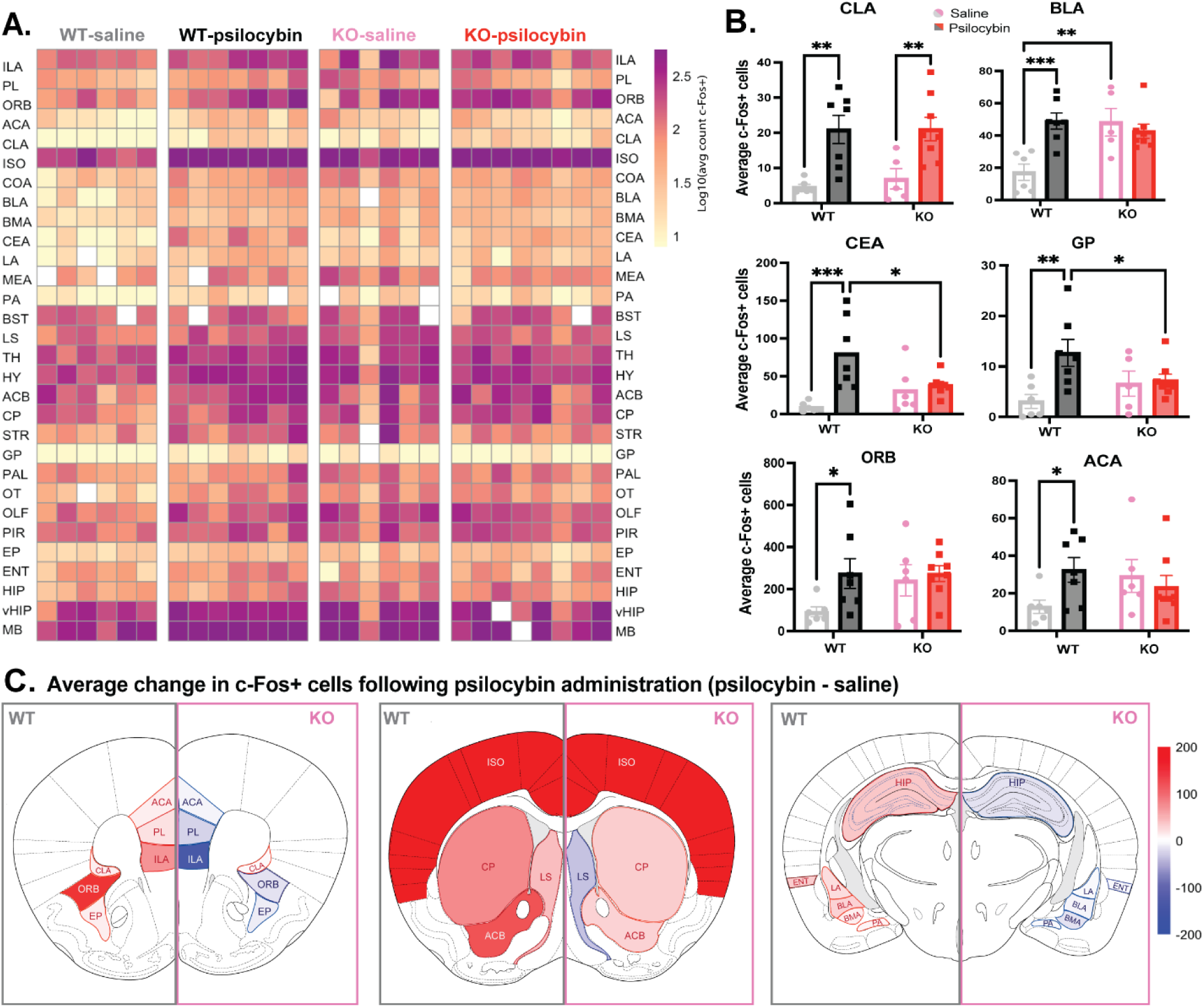
Psilocybin-induced changes in neural activity are dependent on 5-HT1BR **(A)** Heatmap shows log-transformed counts of c-Fos+ cells for individual mice (columns) across brain regions (rows) for control and 5-HT1BR knockout (KO) mice treated with saline or psilocybin. White squares indicate missing data points. **(B)** Group means of c-Fos+ cells in six representative brain regions following psilocybin or saline treatment. **(C)** Percent change in c-Fos+ cells following psilocybin administration is shown in three diagrams of coronal sections of the brain (at bregma +1.94mm, +0.98mm, and -2.18mm) in control (left hemisphere of each diagram) and 5-HT1BR knockout (right hemisphere of each diagram) mice. Psilocybin leads to both increases (shown in red) and decreases (blue) in c-Fos+ cell counts, with notable differences between genotypes, calculated as relative changes in c-Fos expression compared to saline-injected controls, with increases colored as red if they were greater than 200%. Data are presented as mean ± SEM. *p < 0.05, **p < 0.01, ***p < 0.001. ILA: infralimbic cortex, PL: prelimbic cortex; ORB: orbitofrontal cortex, ACA: anterior cingulate cortex, CLA: claustrum, ISO: isocortex, COA: cortical amygdala, BLA: basolateral amygdala, BMA: basomedial amygdala, CEA: central amygdala, LA: lateral amygdala, MEA: medial amygdala, PA: posterior amygdala nucleus, BST: bed nucleus of the stria terminalis, LS: lateral septum, TH: thalamus, HY: hypothalamus, ACB: nucleus accumbens, CP: caudate putamen, STR: dorsal striatum, GP: globus pallidus, PAL: pallidum, OT: olfactory tubercle, OLF: olfactory areas, PIR: piriform cortex, EP: endopiriform nucleus, ENT: entorhinal area, dHIP: dorsal hippocampus, vHIP: ventral hippocampus, MB: midbrain

### 5-HT1BR may influence some of the acute behavioral effects of psilocybin

To investigate the functional effect of 5-HT1BR activation on the acute behavioral response to psilocybin, we measured the head twitch and locomotor response to psilocybin in male and female control and 5-HT1B KO mice. Mice treated with psilocybin demonstrated significantly more head twitch responses than mice treated with saline (Fig 2A; main effect of psilocybin: F (1, 28) = 312.3). However, there was no significant difference in the number of head twitches between genotypes (main effect of genotype: F (1, 28) = 0.4982), indicating that psilocybin robustly induces head twitches in mice independent of 5-HT1BR. This is consistent with previous research implicating 5-HT2AR activation in this acute response to psilocybin (55,56), and demonstration of normal levels of 5-HT2AR expression in mice lacking 5-HT1BRs (57). We also analyzed the locomotor response to psilocybin in mice lacking 5-HT1BR expression, and found that unlike the head twitch response, 5-HT1BR may be involved in the acute hypolocomotor effects (Fig 2B). Specifically, psilocybin significantly decreased locomotion in the 30min following administration (F(1, 48)=8.18, p<0.0063), without effects on velocity (Fig S3A). This effect seemed attenuated in mice lacking 5-HT1BR, however this interaction didn’t reach statistical significance (genotype by psilocybin interaction F(5, 240)=2.00, p=0.0791). Using AUC values for planned comparisons within genotype (Fig 2C), there were significant reductions in locomotor activity following psilocybin in control (t_28_=2.92, p=0.0068), but not in 5-HT1BR KO mice (t_20_=1.26, p=0.16). Overall, these data show that 5-HT1BR could partially mediate some acute behavioral responses to psilocybin, though not the classic head twitch response.

**Figure 2:**
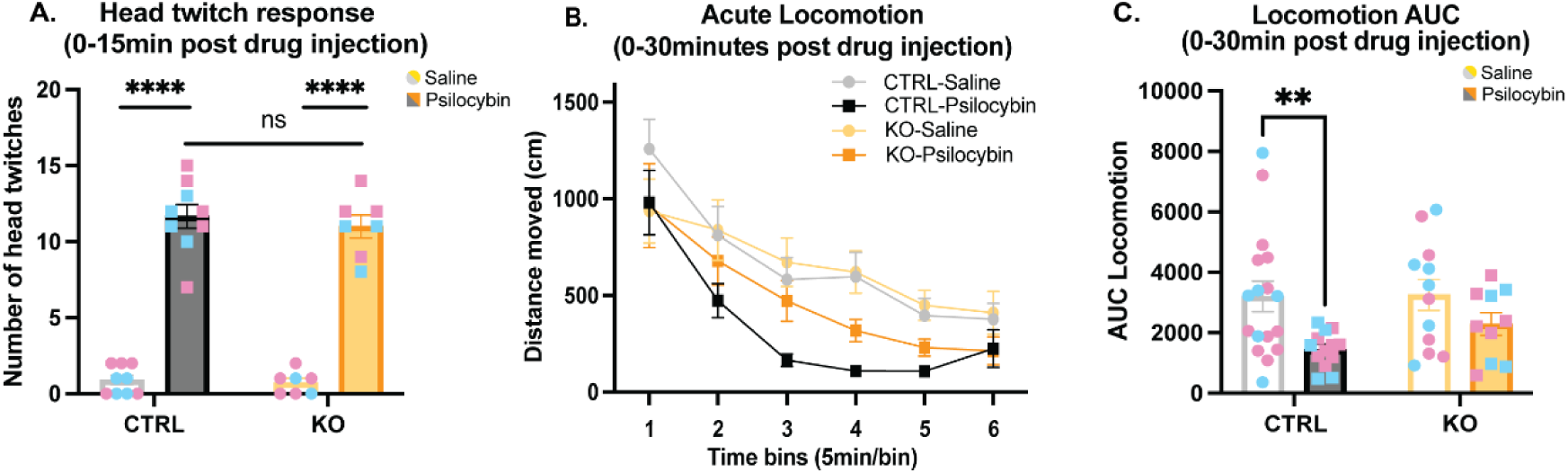
**5-HT1BR may be involved in some of the acute effects of psilocybin** (A) Head twitch response is shown for 5-HT1BR KO mice and littermate controls over the 15-minute period following administration of saline or psilocybin. Psilocybin treatment significantly increased the number of head twitches in both controls and KO mice compared to saline-treated controls with no significant difference between genotypes. (B) Locomotor activity is shown for 5-HT1BR KO mice and littermate controls mice over the 30 minutes following administration of saline or psilocybin. (C) Area under the curve of the locomotor activity over the 30 min following treatment summarizes the locomotor effects of psilocybin. Individual animals are shown as circles (saline) or squares (psilocybin) and colored by sex (pink for females, blue for males). Group data is represented as mean ± SEM. *, p < 0.05; ****p < 0.0001.

### 5-HT1BR is required for antidepressant-like effects of psilocybin in female mice treated with corticosterone

Given that there are long lasting behavioral effects of a single administration of psilocybin in humans, we next wanted to examine the role of the 5-HT1BR in the persisting behavioral effects in mice. We used two paradigms to model stress-induced anhedonia and anxiety-like behaviors. First, we administered chronic corticosterone in the drinking water for four weeks to induce a stress-phenotype to measure antidepressant-like effects of psilocybin (Fig 3A), which did not influence the acute responses to psilocybin (Fig S3B-C). Interestingly, we tested mice lacking 5-HT1BR expression and littermate controls in a series of behavioral tests that evaluated anhedonia and anxiety-like behaviors, performed 24-72h after a single i.p. injection of either vehicle or 5mg/kg psilocybin. In female mice, psilocybin significantly increased hedonic responding in the gustometer as measured by increased licking for sucrose (Fig 3B; treatment by sucrose concentration: F(2,384) = 5.71, p = 0.004). There was also a significant effect of genotype on licking across sucrose concentrations (F(1, 384) = 9.12, p = 0.003) which is consistent with the baseline differences between genotypes that we have previously reported (58). Though the interaction of genotype and treatment did not reach statistical significance (F(2, 384)=2.53, p=0.081), because there were baseline differences, we compared the effect of psilocybin within each genotype. While control mice showed a significant effect of treatment on sucrose consumption (Fig 3B; F (2, 42) =6.09, p=0.005), there was no effect in female mice lacking 5-HT1BR (F(2, 30) = 0.73, p=0.489). Looking specifically at consumption of the highest concentration of sucrose in control mice (Fig 3C; main effect of treatment F(2, 72)=5.66, p=0.005), psilocybin increases consumption compared to saline treated mice (p=0.012), restoring it back to non-cort baseline levels (p=0.266). In 5-HT1BR KO mice, although cort decreased sucrose consumption (no cort/saline vs cort/saline: p=0.030), psilocybin did not significantly increase the consumption (cort/saline vs cort/psilocybin: p=0.549). Overall, these data suggest that psilocybin may influence anhedonia in mice through actions at the 5-HT1BR.

**Figure 3:**
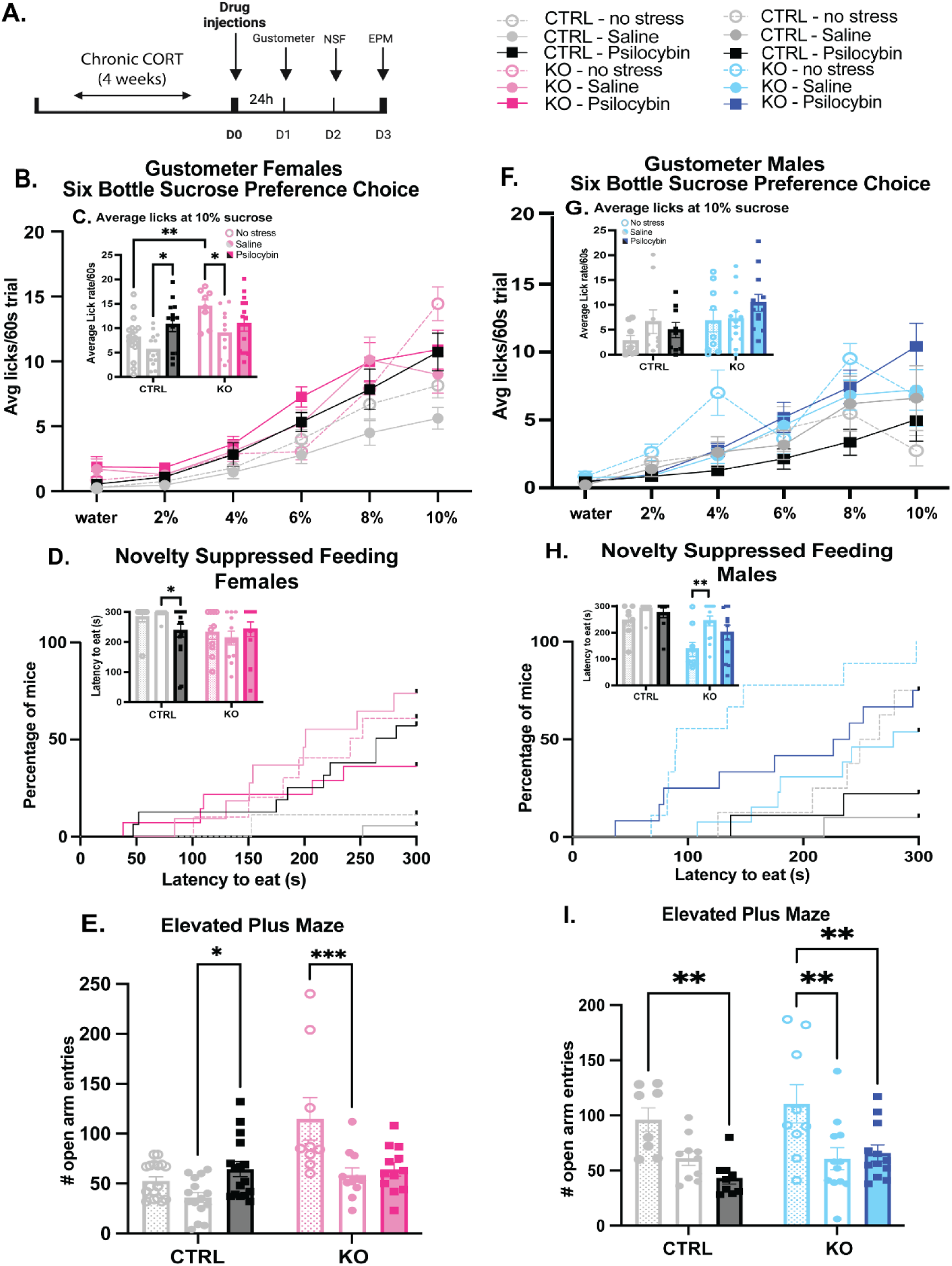
5-HT1BR is required for behavioral effects of psilocybin in female mice. (A) Experimental timeline is shown for studies testing the post-acute behavioral effects of psilocybin. **(B)** Gustometer data shows the number of licks in females across 6 concentrations of sucrose in female control and 5-HT1BR KO mice, following chronic cort with saline or psilocybin treatment, and for the no stress control condition. **(C)** Inset shows the number of licks at the 10% sucrose concentration in females. **(D)** Kaplan-Meier survival curves represent the cumulative proportion of females eating over time in the NSF for controls (shades of black) and 5-HT1BR KOs (shades of pink). Inset shows individual data points and average group latency to eat. **(E)** Open arm entries are shown for females in the EPM for controls and KOs over the three treatment groups. **(F)** Gustometer data shows the number of licks in males across 6 concentrations of sucrose in female control and 5-HT1BR KO mice, following chronic cort with saline or psilocybin treatment, and for the no stress control condition. **(G)** Inset shows the number of licks at the 10% sucrose concentration in males. **(H)** Kaplan-Meier survival curves represent the cumulative proportion of males eating over time in the NSF for controls (shades of black) and 5-HT1BR KOs (shades of pink). Inset shows individual data points and average group latency to eat. **(I)** Open arm entries are shown for males in the EPM for controls and KOs over the three treatment groups. Data points on bar graphs represent individual mice and group averages are shown as mean ± SEM. *p<0.05, **p<0.005, ***p<0.001

Expression of 5-HT1BR also significantly modulated the anxiolytic response to psilocybin in female mice. First, we measured anxiety-like behavior in the conflict-based novelty-suppressed feeding (NSF) assay 48h following psilocybin administration. Interestingly, there was an anxiolytic response to psilocybin that was significantly blunted in the absence of 5-HT1BR expression (genotype by drug interaction: F(2,72) = 2.99, p = 0.049). Specifically, control mice injected with psilocybin showed reduced latency to eat compared to saline-treated mice (Fig. 3D, χ^2^ = 9.62, p = 0.002), while we saw no significant effect in the 5-HT1BR KO mice (Fig. 3D, χ^2^ = 1.85, p = 0.174). There was also a main effect of genotype independent of treatment (F(1,72)=7.29, p=0.009), suggesting that there are again baseline effects of brain-wide, life-long absence of 5-HT1BR expression on anxiety-like behavior. There were no significant effects of genotype or drug treatment in the home cage condition suggesting that the effects of psilocybin were on anxiety behavior rather than feeding behavior (Fig S4A). We also tested mice in the elevated plus maze (EPM), as a second measure of anxiety-like behavior, 72h following psilocybin administration. We again found that psilocybin-induced decreases in anxiety behavior that were dependent on 5-HT1BR expression (Fig. 3E; main effect of drug by genotype (F(1, 51) = 4.080, p = 0.0487). Psilocybin administration significantly decreased anxiety in female control mice, as measured by increased entries in open arms (p=0.01), but not in mice lacking 5-HT1BR (p = 0.732). This was also somewhat reflected in the time spent in open arms of the EPM (Fig S4B), and was not due to any persisting changes in overall locomotion (Fig S4C). Overall, convergent evidence from two conflict-based assays shows that psilocybin’s persisting anxiolytic response requires 5-HT1BR. Overall, these behavioral results indicate that in female mice, psilocybin produces an antidepressant-like phenotype in the days following administration which we measured as decreased anhedonia in the gustometer and decreased anxiety-like behaviors in the NSF and EPM. Interestingly, these behavioral effects were attenuated in the absence of 5-HT1BR expression, suggesting that the antidepressant-like effects of psilocybin are dependent on 5-HT1BR following chronic corticosterone administration in female mice.

Interestingly, in male mice, we were unable to find any behavioral effect of psilocybin following chronic corticosterone treatment, which is consistent with previous findings (27). Specifically, psilocybin had no effect on hedonic responding in the gustometer as measured by licking for various sucrose concentrations (Fig 3F; F(2, 230.4)=2.14, p=0.12). Similar to the females, we again saw an effect of genotype on licking over the sucrose concentrations in males (F(1, 294)=11.36, p<0.001), as previously reported (58), however there was no significant effect of treatment (F2, 294)=2.37, p=0.1).

Analyzed within each genotype, there was no effect of psilocybin in either 5-HT1BR KOs or controls (F(2,23)=0.60, p=0.55 for controls, F(2, 31)=1.1, p=0.34 for KOs). Similar patterns were seen when looking at consumption of the highest concentration of sucrose (Fig 3G). There was a significant genotype effect, but no effect of treatment (F(1, 54)=5.3, p=0.028 for genotype, F(2, 54)=1.3, p=0.28 for treatment).

We saw a similar trend for males in two tests of anxiety-like behavior – there were effects of cort treatment and of genotype, but no significant effects of psilocybin on these measures. In the NSF test (Figs 3H and S4D) 5-HT1BR KOs showed reduced anxiety-like behavior (F(1, 55)=18.12, p<0.0001), but again there were no effects of psilocybin, despite cort treatment increasing anxiety-like behavior (main effect of treatment: F(2,55)=5.92, p=0.005, veh/saline vs cort/saline: χ_1_ = 6.72, p=0.006 for controls, χ_1_ = 11.13, p=0.001 for KOs; cort/saline vs cort/psilocybin: χ_1_ = 0.50, p=0.48 for controls, χ_1_ = 1.66, 0.24 for KOs). In the EPM (Figs 3I, 4SE-F), there were no effects of genotype on the number of entries into the open arms in males (F(1,54)=2.22, p=0.14). There was a significant effect of treatment (F(2, 54)=12.64, p<0.0001), though this effect was driven by the effect of cort (veh/saline vs cort/saline: p=0.0007), rather than psilocybin (cort/saline vs cort/psilocybin: p=0.57). Overall, although genotype and cort had some behavioral effects in males, psilocybin consistently did not influence anhedonia and anxiety-like behavior.

**Figure 4:**
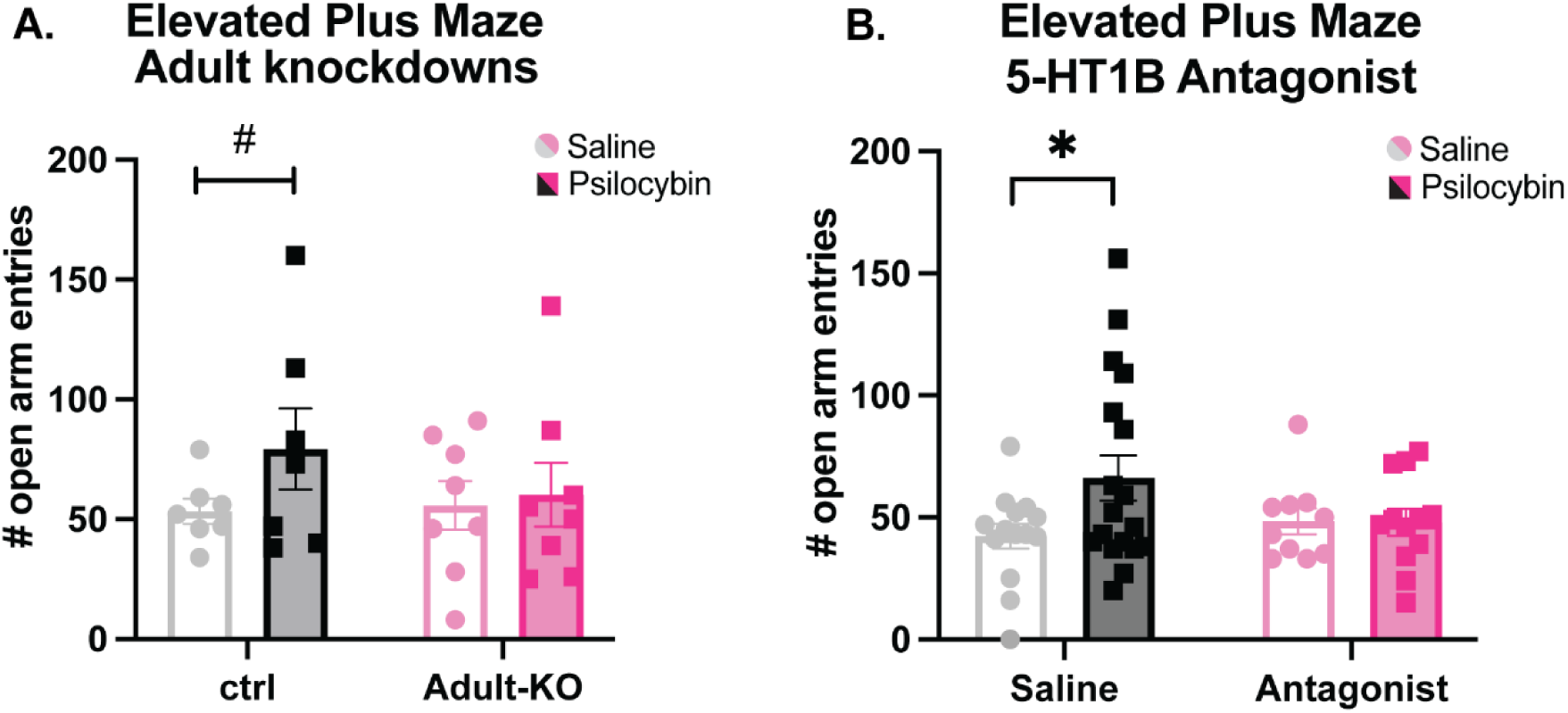
(A) There was no significant effect of psilocybin in mice with adult knockout of 5-HT1BR expression, however the effect of psilocybin in control mice did not reach statistical significance in this cohort either. **(B)** Psilocybin significantly increased entries into the open arms of the EPM in saline-treated controls mice, suggesting reduced anxiety. However, pre-treatment with the 5-HT1BR antagonist GR127935 blocked this effect, indicating the importance of 5-HT1BR activation during the acute phase of psilocybin for its anxiolytic effects. #p=0.1; *p < 0.05

Acknowledging potential compensatory effects of the absence of 5-HT1BR throughout development, we used two approaches to address this alternative interpretation of the psilocybin effect in females. First, we used an adult-knockdown approach, in which mice were reared with normal expression of 5-HT1BRs through PN60 via administration of doxycycline to block tTS-mediated knockdown (as previously reported and validated (58)). Following withdrawal of doxycycline as adults, mice lacking 5-HT1BR expression were tested for the persisting behavioral effects of psilocybin during chronic corticosterone administration in the elevated plus maze (Fig 4A). Though limited in number, a planned comparison shows no significant effect of psilocybin in adult knockdown mice (t_14_=0.26, p=0.80). Though the interpretation of this result is limited by an effect in controls that was only trending toward significance (t12=1.461, p=0.11), a lack of effect in the adult knockdown suggests that any developmental compensation was likely not responsible for the lack of effects seen in the whole-brain whole-life knockouts. This effect was mirrored in the time spent in the open arms (Fig S5A), and was not due to differences in locomotion (Fig S5B). Additionally, to investigate if the behavioral effects in the constitutive 5-HT1BR KO were due to activation of the 5-HT1BR during the acute phase of psilocybin, and to attempt to circumvent genotype differences seen in some of the behavioral tests, we performed another experiment in which we pharmacologically blocked 5-HT1BR with pre-treatment with a 5-HT1B/1D receptor antagonist, GR127935, during psilocybin administration in control mice. Psilocybin again decreased anxiety-like behavior as measured by increased open-arm entries as a function of 5-HT1BR activation (Fig. 4B, pretreatment x psilocybin interaction: F(1, 48)=4.30, p=0.044). Specifically, in the absence of any pretreatment, psilocybin increased open arm entries (p=0.0084), however when pre-treated with GR127935, the effect of psilocybin was blocked (p=0.89). Measures of open arm duration or locomotion were not significantly influenced by treatment (Fig S5C-D). Overall, these results suggests that blockade of 5-HT1BR during the acute psilocybin response may be sufficient to block the post-acute anxiolytic effects of psilocybin seen in the EPM.

### 5-HT1BR mediates the anxiolytic effects of psilocybin in a behavioral despair model in both female and male mice

We used a second stress paradigm to confirm the results seen in the female mice in the chronic cort model, and because the chronic cort model was not effective to measure the persisting behavioral effect of psilocybin in male mice. Using a chronic behavioral despair paradigm implemented through repeated forced swim stress, we examined the effect of the absence of 5-HT1BR on the behavioral effects of psilocybin in mice (Fig 5A). Interestingly, in this paradigm we were able to elicit an effect of psilocybin in both sexes. There were significant effects of treatment on consumption across the sucrose concentrations (Fig 5B; treatment x concentration: F(2, 538)=3.63, p=0.027). Given that we saw a baseline genotype effect again (genotype x concentration: F(1, 538)=10.29, p=0.001; genotype x treatment x concentration F(2, 538)=2.41, p=0.09), we analyzed the treatment effects within each genotype separately. In control mice, there was a significant effect of treatment on sucrose consumption (treatment x concentration: F(10, 330)=2.87, p=0.002), which was largely driven by a increase in consumption of 10% sucrose following psilocybin compared to saline treatment (Fig 5C; p=0.009). Though there were no significant sex differences (F(1,488)=0.86, p=0.35) and both sexes showed similar trends, the effects of psilocybin in females may be greater (Figs S6A-D). On the other hand, there was no effect of treatment in 5-HT1BR KO mice (F(10, 190)=0.75, p=0.68), suggesting that stress and/or psilocybin don’t significantly influence hedonic responding in the absence of 5-HT1BR expression in mice.

**Figure 5.**
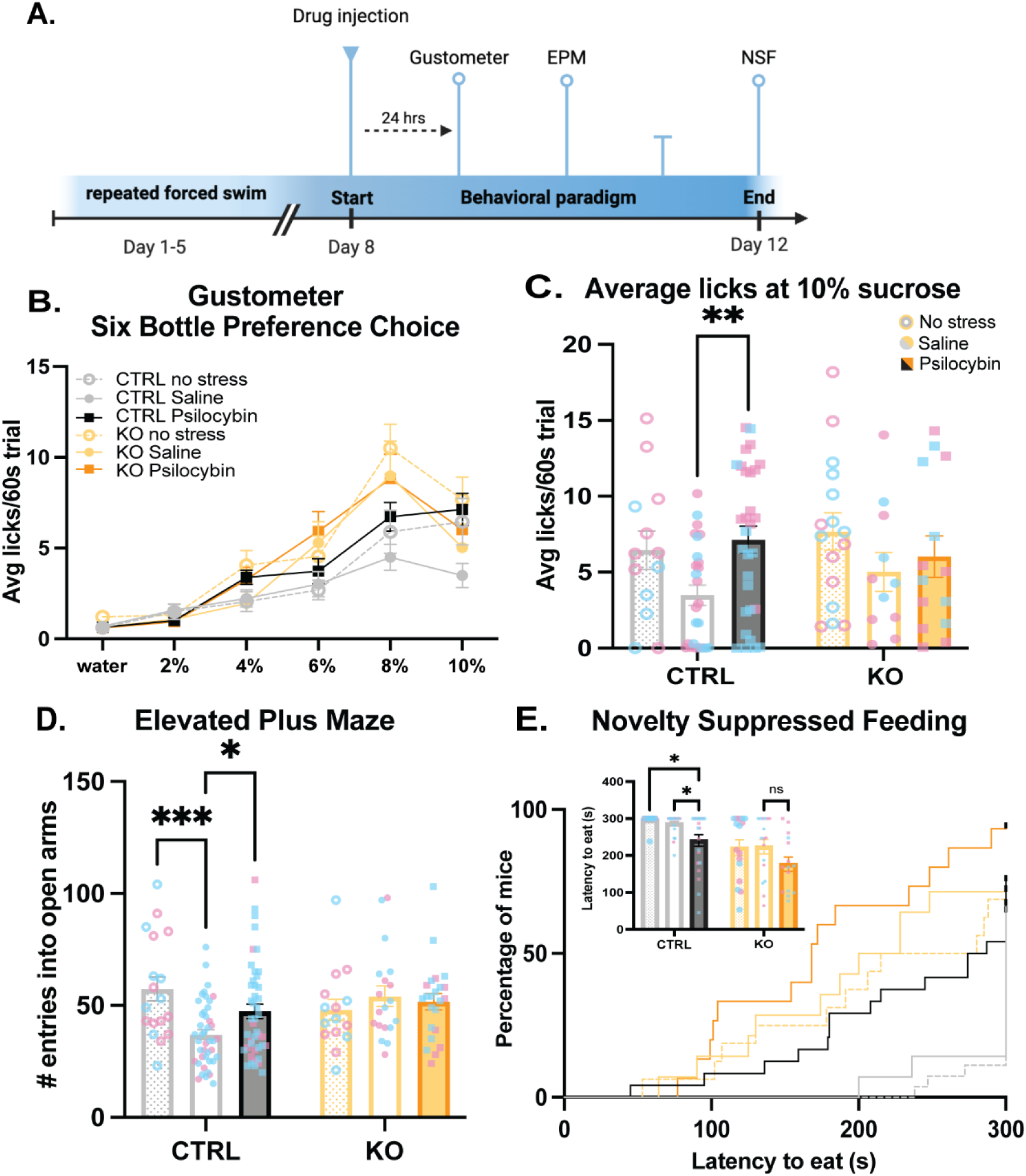
The effects of Psilocybin on behavior in a behavioral despair model are mediated by 5-HT1BR expression in both male and female mice. **(A)** Experimental timeline is shown for testing the post-acute behavioral effects of psilocybin following forced swim stress exposure. **(B)** Gustometer data shows the number of licks across 6 concentrations of sucrose in control and 5-HT1BR KO mice, following forced swim stress with saline or psilocybin treatment, and for the no stress control condition. **(C)** The number of licks at the 10% sucrose concentration is shown for control and 5-HT1BR KO mice in the three treatment conditions. **(D)** Open arm entries are shown in the EPM for both sexes in control and 5-HT1BR KO mice in three treatment groups. **(E)** Kaplan-Meier survival curves represent the cumulative proportion of mice eating over time in the NSF for controls (shades of black) and 5-HT1BR KOs (shades of yellow). Inset shows individual data points and average group latency to eat. Data points on bar graphs represent individual mice, with females colored pink and males colored blue. Group averages are shown as mean ± SEM. *p<0.05, **p<0.005, ***p<0.001

In the EPM, there was a significant effect of genotype on the anxiolytic effect of psilocybin (Fig 5D; interaction of genotype and treatment: F(2, 155)=4.22, p=0.016). In control mice, stress decreased entries into the open arms (p<0.0005), and psilocybin rescued this effect (p=0.03). Interestingly, in the mice lacking 5-HT1BR expression, psilocybin had no effect on entries into the open arms (p=0.99), though stress also had no effect in this strain (p=0.84). This pattern was replicated in the duration of time spent in open arms of the EPM (Fig S6E-G). There were also no effects on locomotion generally in the EPM (Fig S6H-J), and no significant effects of sex on the EPM behavior (genotype x treatment x sex: F(2, 155)=0.146, p=0.86).

In a second measure of anxiety behavior in the NSF assay, psilocybin also decreased anxiety-like behavior as measured by latency to eat (Fig 5E, X_5_=31.40, p<0.0001). There were again some baseline differences between genotypes, but importantly within control mice, psilocybin decreased the latency to eat (stress/saline vs stress/psilocybin: X_1_=6.41, p=0.011), but had no significant effect in 5-HT1BR KO mice (X_1_=1.70, p=0.19). There were again no effects of sex on the treatment effects (sex by treatment: F(2, 104)=0.14, 0.87). There were no effects of genotype or psilocybin on latency to eat in the homecage suggesting that hunger was not a cause of the differences in behavior (Fig S6K-M).

### Neural circuits impacted by psilocybin are influenced by 5-HT1BR expression

To identify the neural circuits through which 5-HT1BR may mediate the effects of psilocybin on behavior, we performed a network analysis on neural activity following psilocybin to probe functional connectivity. Returning to the c-Fos data (shown in Fig 1), we generated pairwise correlation matrices of brain region activity as measured by number of c-Fos+ cells to determine how the activity of brain regions vary together. A difference matrix for each genotype, representing the change in correlated activity following psilocybin administration, illustrates how the functional connectivity throughout the brain is changed by psilocybin and differs based on 5-HT1BR expression (Fig. 6A). Comparing these two correlation matrices reveals statistically distinct patterns between control and 5-HT1BR KO mice (Fig 6A; rho = -0.01, p = 0.80). Control mice showed more desynchrony following psilocybin compared to saline (t = -5.336 with p<0.0001).

**Figure 6:**
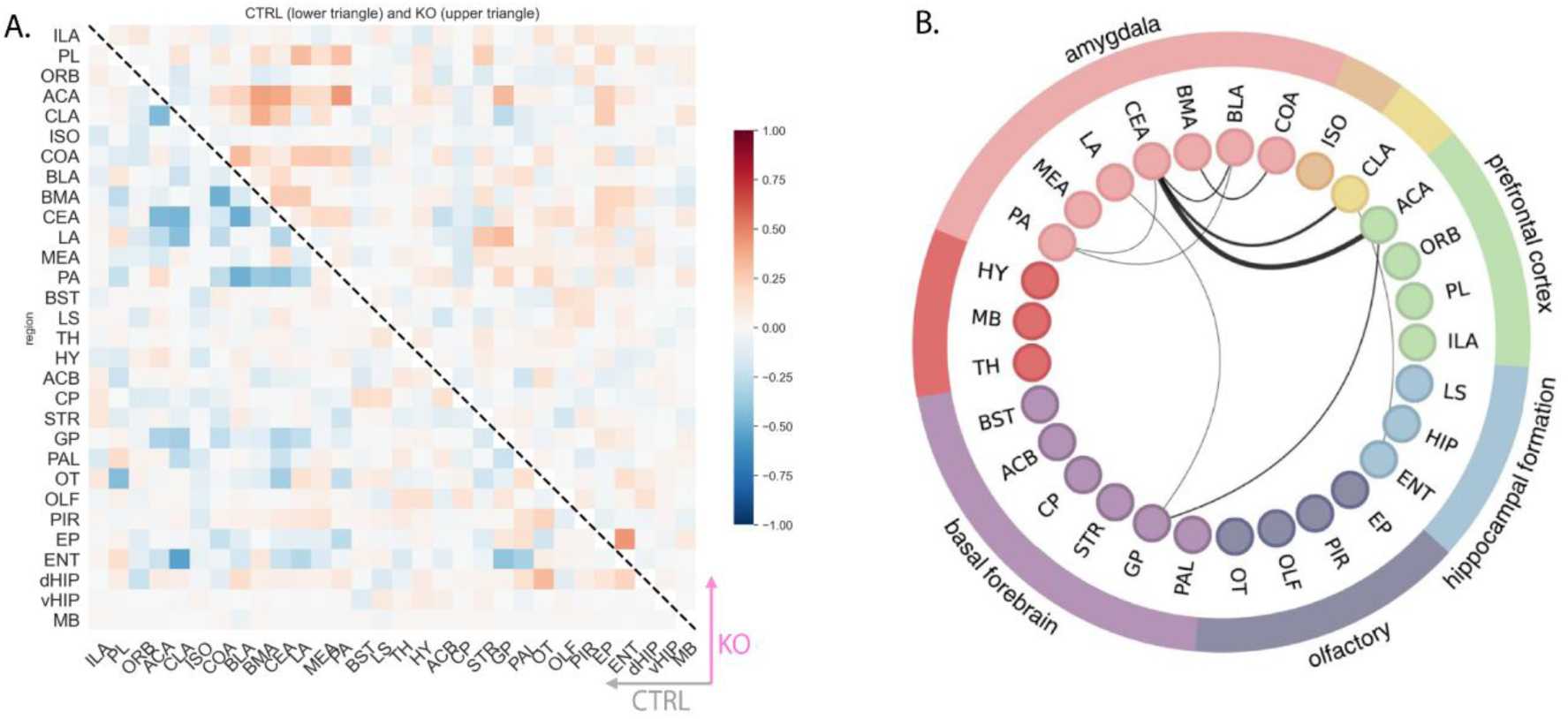
**5-HT1BR expression influences select neural circuits influenced by psilocybin** (A) The difference correlation matrix shows psilocybin-induced alterations in functional connectivity between control (lower triangle) and 5-HT1BR KO mice (upper triangle). The matrix represents the pairwise correlation differences across 30 regions, illustrating desynchrony patterns between the two genotypes, with control mice exhibiting more desynchrony following psilocybin than 5-HT1BR KO mice do. (B) Circular network plot highlighting psilocybin-responsive connections most influenced by 5-HT1BR expression. Significant changes in correlated activity were observed between the CEA and the ACA (p = 0.05). Expanding the confidence interval to 90%, 85%, and 80% revealed additional differences in connectivity in CEA-CLA, ACA-GP, CEA-COA, CEA-BLA, CLA-ENT, GP-LA, PA-CEA, PA-BLA. Line width corresponds to the confidence interval, with the thickest edge representing 95% CI.

Interestingly, this is reversed in mice lacking 5-HT1BR (t=+5.083 p < 0.0001 for KOs), suggesting that neural activity is more positively correlated across regions following psilocybin treatment. We then examined which psilocybin-responsive networks were most influenced by 5-HT1BR expression and found significant effects of 5-HT1BR on correlated activity between the central amygdala and the anterior cingulate cortex (z=2.00, p=0.05). Widening the confidence interval, we found suggestive differences in the correlated activity of the central amygdala with the claustrum, amongst amygdalar regions, and between the globus pallidum and lateral amygdala and anterior cingulate cortex (Fig 6B). Overall, this network analysis shows that 5-HT1BR expression significantly influences neural circuits that are relevant to the effects of psilocybin on cognition and mood. Furthermore, it points to potential neural circuits through which 5-HT1BR may influence some of the persisting effects of psilocybin.

## Discussion

Our results highlight a critical role for 5-HT1BR in the response to psilocybin in mice. We found that the absence of this non-hallucinogenic receptor impacts the neural response to psilocybin, as well as the acute and persisting behavioral effects. Using c-Fos as a marker of neural activity, we find increased activity in many regions throughout the brain including in sensory and prefrontal cortices, claustrum, amygdala, hypothalamus, and basal ganglia, which is consistent with previous reports (22,59–61). Interestingly, we saw that mice lacking expression of 5-HT1BR have notable differences in the pattern of neural activity following psilocybin administration, including in prefrontal, hippocampal, and amygdalar areas, all brain regions that express high levels of 5-HT1BR. Functionally, a lack of 5-HT1BR had no effect on the head twitch response, though it did attenuate the acute hypolocomotor effects of psilocybin. Importantly, there were also effects of 5-HT1BR on the persisting behavioral effects of psilocybin seen by the amelioration of stress-induced anhedonia and anxiety-like behaviors. Overall, this shows that 5-HT1BR expression is involved in mediating some the neural and behavioral effects of psilocybin.

Though we find that 5-HT1BR is required for the persisting effects of psilocybin, our data do not address its sufficiency. The profound neural and behavioral effects of psilocybin are likely supported by its polypharmacology at multiple serotonin receptors (32).

Although there is some evidence questioning the necessity of the 5-HT2AR in the behavioral effects of psilocybin in mice (23,26), there are mixed findings on this topic with other reports showing that 5-HT2AR expression is necessary for the anti-depressant effects of psilocybin (22). Tangential to this debate, our results show that acute activation of the 5-HT2A receptor by psilocybin is insufficient to rescue stress-induced behaviors in the absence of the 5-HT1BR.

There are many lines of evidence linking the 5-HT1BR to antidepressant function as well as to the behavioral and neural response to hallucinogens (62,63). As an inhibitory G_i/o_-coupled receptor, its activation reduces calcium influx in axon terminals of serotonin (autoreceptors) and non-serotonin (heteroreceptors) neurons decreasing neurotransmitter release (64–66). Past research shows that 5-HT1BR agonists promote antidepressant-like effects and can increase the rewarding effects of cocaine and social interaction (67–70). 5-HT1BR also interacts with p11 to regulate depression states (71,72). Additionally, 5-HT1BR is necessary for the behavioral and hippocampal plasticity effects of fluoxetine (33). In addition to mediating depressive phenotypes, there is also evidence implicating the 5-HT1BR in antidepressant response to ketamine (39,42), and in other behavioral effects of MDMA (38,40,41).

In keeping with our previous work showing that the 5-HT1BR has a known role in mediating reward and anxiety behavior (36,43,58), we do find baseline differences in some of the behaviors we tested in 5-HT1BR KO mice. Additionally, there were also behavioral assays in which mice lacking 5-HT1BR seemed to show more resilience to chronic corticosterone treatment, particularly in the gustometer measure of anhedonia. These differences may arise from compensatory changes due to the absence of 5-HT1BR throughout development. Alternatively, these baseline differences could be a reflection of the important role of 5-HT1BR in mediating these phenotypes in adulthood. However, the baseline differences leave open the limitation of potential behavioral and mechanistic ceiling effects which could have obscured the effects of psilocybin in this strain. This would be possible either through maxing out the range of the behavioral assays and/or through saturation of the 5-HT1BR-dependent mechanism through which psilocybin has its effects. We first addressed these limitations by using at least one behavioral assay with no baseline differences, namely the elevated plus maze.

Additionally, we tested mice with adult knockdown of the receptor to eliminate any developmental compensation. Finally, we used a pharmacological blockade of 5-HT1BR by administering an antagonist prior to psilocybin administration. This not only allowed us to circumvent any baseline differences, but also helped identify that 5-HT1BR is involved during the acute drug effect.

Consistent with past research, we find some sex differences in our behavioral effects of psilocybin. We found that the post-acute behavioral effects of psilocybin on anhedonia and anxiety-like behavior were influenced by an interaction of stressor-type and sex.

Specifically, psilocybin was able to rescue the anhedonia and anxiety-like behavior in females exposed to chronic corticosterone treatment, however, this effect was not seen in males. This sex difference may have emerged from a differential response to the stress paradigm rather than a differential response to psilocybin, particularly due to a more robust effect of the corticosterone treatment on the behavior of males. This is consistent with past work showing that there are sex-specific effects of chronic corticosterone administration in the drinking water (73,74). On the other hand, using a behavioral despair model via repeated forced swim stress, psilocybin was able to rescue stress-induced behavioral phenotypes in both male and female mice. This suggests that different stress modalities may induce sex-dependent alterations by engaging different neural circuits that are amenable to varied pharmacological interventions. For example, recent work shows that psilocybin differentially influences hypothalamic reactivity to aversive stimuli and reduces ethanol consumption specifically in male, but not female rodents (59,75). On the other hand, females display more head twitch behavior to the psychedelic 1-(2,5-dimethoxy-4-iodophenyl)-2-aminopropane (DOI) and less of an augmentation of pre-pulse inhibition in males, in a strain dependent manner (76,77).

Ultimately, there appears to be a substantial impact of sex on the effect of psilocybin on behavior, which are dependent on strain and paradigm, reinforcing the importance of examining sex differences in future studies.

While the 5-HT2AR plays a central role in psilocybin’s acute psychedelic effects, its therapeutic potential in the context of stress-induced depression may be limited without the support of other serotonin signaling pathways, including those acting via 5-HT1B. The absence of 5-HT1BR signaling may prevent psilocybin from achieving its full therapeutic potential as the feedback inhibition of serotonin release is disrupted, leading to overactivation of serotonergic circuits, and limiting the ability to modulate neural mechanisms effectively. This could result in a mechanistic ceiling of 5-HT2A, essentially maxing out the 5-HT2A-dependent effects on anhedonia and anxiety-like behaviors. This hypothesis arises in part from studies showing increased 5-HT2AR expression following chronic exposure to stress (78–80). However, neither stress nor the knock-out model seem to have affected acute 5-HT2AR activation in our study, as measured by head twitch responding, and past studies showing no effect of 5-HT1BR KO on 5-HT2AR binding (57). Additionally, although our hypothesis of 5-HT1BR involvement is via direct binding of psilocin to 5-HT1BRs on neural circuits implicated in anxiety and depressive behaviors, it is also possible that these effects may be indirect. For example, 5-HT1BR could influence the hypothermic response to psilocybin which could then influence the antidepressant-like behavioral response (81–83). Overall, in combination with other work, our results highlight that psilocybin’s persisting effects on behavior may depend on an optimal balance of serotonin tone acting across different serotonin receptor subtypes potentially across a number of physiological systems.

Our studies begin to point to the anterior cingulate cortex, the claustrum, and the amygdala as potential targets of psilocybin which are modulated by 5-HT1BR expression. Some preclinical studies have already highlighted the amygdala, especially the basolateral and central nucleus of the amygdala, as important loci for the therapeutic effects of psilocybin (84–87). Given the role of the amygdala in mediating the effects of psilocybin on threat responding (88), it is possible that 5-HT1BR expression on amygdalar terminals underlies our reported behavioral effects. We also found that the 5-HT1BR-effects on functional connectivity seemed to involve the amygdala as a node given its high number of connections with other brain regions that were modulated by 5-HT1BR expression. Our current studies are focusing on region-specific knockdown of 5-HT1BRs in the amygdala to test their functional role in modulating the effects of psilocybin on emotional reactivity. Projection specific studies will also investigate if prefrontal-amygdala networks are involved, and how psilocybin may influence these connections via 5-HT1BR since serotonin is known to gate cortical glutamatergic inputs into the BLA via 5-HT1B receptors (89). Understanding how psilocybin affects interregional connectivity within these networks via its actions on 5-HT1BR activation will help identify the mechanisms through which psilocybin exerts its antidepressant effects.

Overall, our work highlights the complexity of the serotoninergic targets involved in the therapeutic efficacy of psilocybin. Specifically, these studies add to our understanding of the importance of non-5-HT2AR targets by implicating 5-HT1BRs in the antidepressant-like effects of psilocybin in mice. These data contribute to the broader understanding of the polypharmacology of psilocybin and highlight the need for multi-receptor approaches to better understand psilocybin’s mechanisms of action. Additionally, our data demonstrate a dissociation of the acute head twitch response from the longer-term effects of psilocybin, suggesting that different mechanisms of actions may underlie the acute psychedelic effect of psilocybin and the persisting behavioral effects. We suggest that psilocybin’s effects are not solely driven by 5-HT2AR activation, and that activation of other serotonin receptors are critical for its therapeutic potential. These results also raise the potential for development of effective non-hallucinogenic pharmacotherapies targeting 5-HT1BR for the treatment of depressive disorders.

