## Supplemental Figures for "The serotonin 1B receptor is required for some of the behavioral effects of psilocybin in mice"

**Supplemental Material**

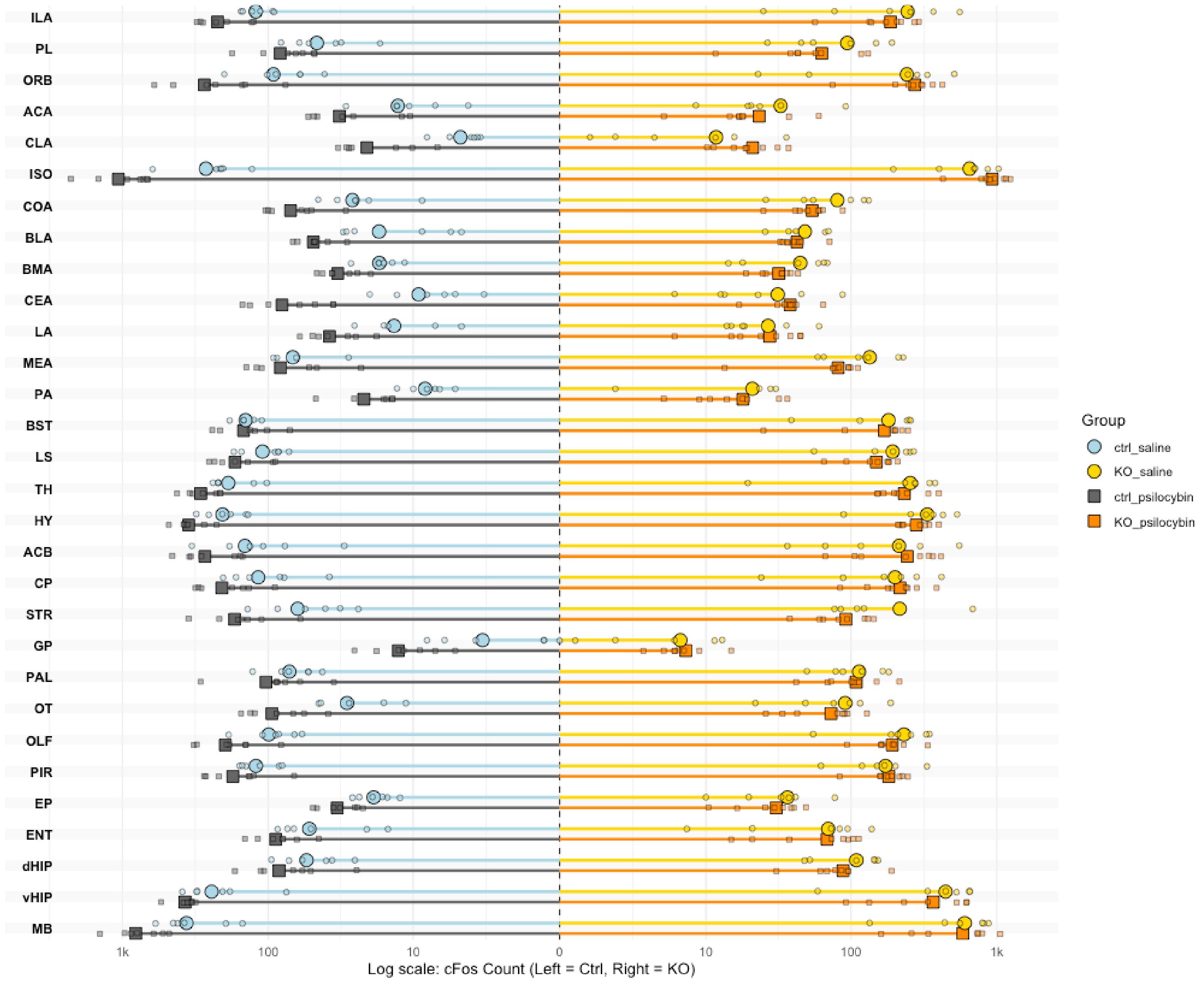


**Supplemental Fig 1: Raw counts of c-Fos+ cells.** Cell counts are shown across all brain regions analyzed in control (left) and 5-HT1BR KO (right) mice treated with psilocybin (grey/orange) or saline (blue/yellow). Individual mice are shown with smaller circles (saline) and squares (psilocybin), and lines ending with larger squares and circles represent group means. ILA: infralimbic cortex, PL: prelimbic cortex; ORB: orbitofrontal cortex, ACA: anterior cingulate cortex, CLA: claustrum, ISO: isocortex, COA: cortical amygdala, BLA: basolateral amygdala, BMA: basomedial amygdala, CEA: central amygdala, LA: lateral amygdala, MEA: medial amygdala, PA: posterior amygdala nucleus, BST: bed nucleus of the stria terminalis, LS: lateral septum, TH: thalamus, HY: hypothalamus, ACB: nucleus accumbens, CP: caudate putamen, STR: dorsal striatum, GP: globus pallidus, PAL: pallidum, OT: olfactory tubercle, OLF: olfactory areas, PIR: piriform cortex, EP: endopiriform nucleus, ENT: entorhinal area, dHIP: dorsal hippocampus, vHIP: ventral hippocampus, MB: midbrain

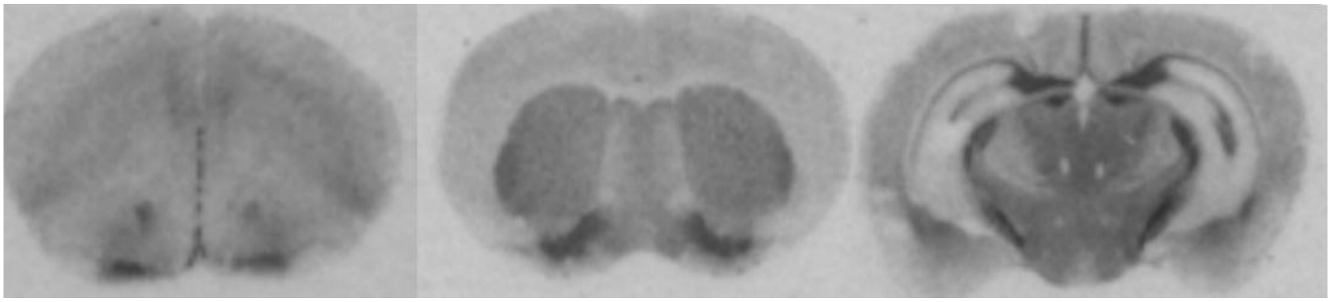

**Supplemental Figure 2: 5-HT1BR protein expression is located in brain regions showing differential c-Fos expression following psilocybin.** Representative images from receptor autoradiography films of control mice processed with [125]I-cyanopindolol for 5-HT1BR localization. Sections were incubated in 70pM of [125]I-cyanopindolol (10µL/10mL buffer; Perkin Elmer), 3µM isoproterenol, 100 nM 8-OH-DPAT, 0.3% BSA, 0.01% ascorbic acid and 10µM pargyline (all from Sigma-Aldrich) for 2h. Slides were then exposed to BioMax MR film (Kodak) for a period of 6-18h, and developed using an X-omat automated developer (Kodak).

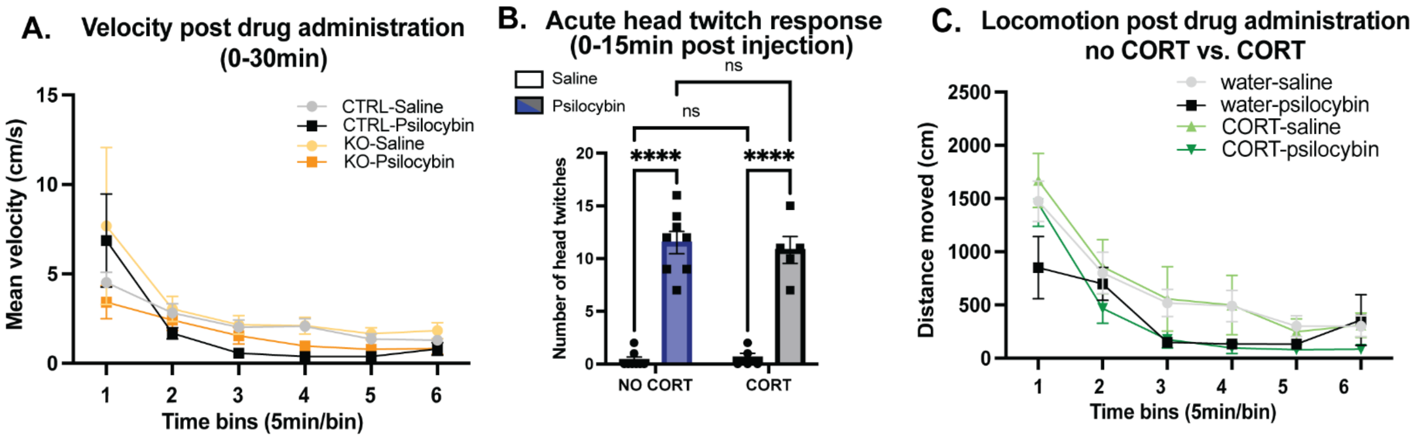


**Supplemental Figure 3: Acute Effects of Psilocybin in Mice. A) Velocity of Control and 5-HT1BR KO mice following acute administration of psilocybin or saline.** There was a significant effect of time on velocity ($F(5,220)=11.59$, $p<0.0001$), but no significant effects of genotype or drug ($F(1,44)=0.35$, $p=0.555$ for genotype, $F(1, 44)=3.68$, $p=0.062$). **B) Effects of cort on head twitch response.** Mice treated with psilocybin exhibited a significantly higher number of head twitches compared to those treated with saline (main effect of psilocybin: $F(1, 22) = 151.1$, $p < 0.0001$), both in the presence and absence of CORT (main effect of CORT: $F(1, 22) = 0.075$, $p=0.787$. The effects were not significantly different between groups receiving CORT versus those not receiving CORT (main interaction of psilocybin x CORT: $F(1, 22) = 0.284$, $p=0.599$). Error bars represent SEM. ns indicates no significant difference, *$p < 0.05$, **$p < 0.01$. **C) Effects of cort on locomotion post-drug administration.** Initial locomotion decreased over time (main effect of time: $F(2.393, 38.29) = 47.82$, $p<0.001$), with psilocybin-treated mice showing lower overall locomotion compared to saline-treated mice (main effect of psilocybin: $F(1, 16) = 5.035$, $p=0.039$), with no difference between groups receiving CORT or regular water (main interaction of drug x CORT: $F(1, 16) = 0.022$, $p=0.883$). Error bars denote SEM.

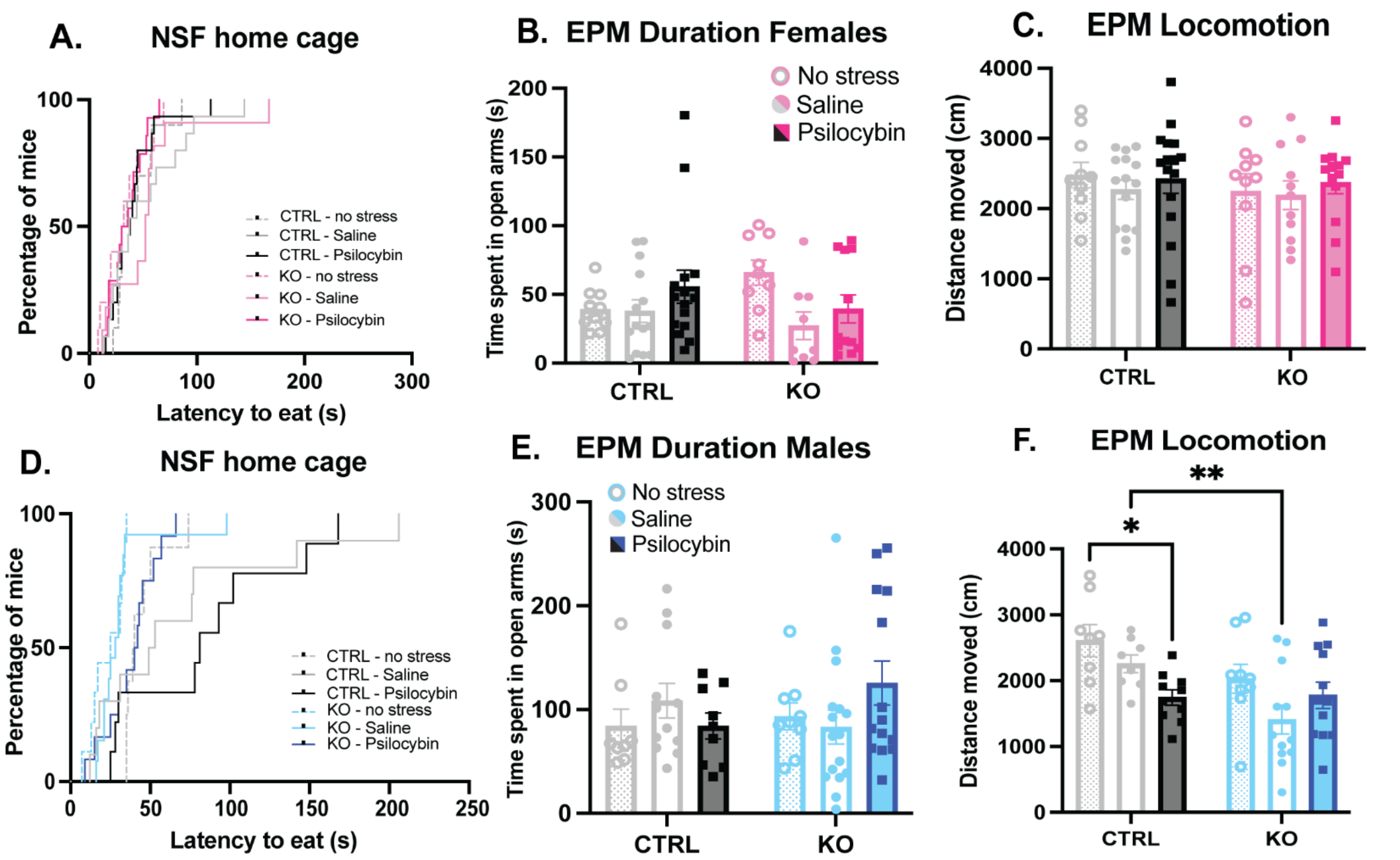


**Supplemental 4: Additional behavioral metrics corresponding to the effect of psilocybin in the cort model in female and male mice shown in Fig 3. (A) Home cage feeding in the NSF in females.** Control females and 5-HT1BR KO females displayed similar eating behavior in the home cage following the NSF, regardless of treatment group (Log-Rank Mantex Cox test, chi square = 2.314, $p = 0.31$). **(B) Time spent in open arms in the EPM in females.** There was a trend towards an interaction of genotype by treatment in time spent in open arms in female mice, mirroring the effects on the number of entries into the open arms (main interaction genotype x treatment: $F(2, 63) = 2.529$, $p = 0.09$). 5-HT1BR KO females showed a significant reduction in time spent in open arms following corticosterone treatment ($F(2,27) = 3.591$, $p = 0.04$), which psilocybin failed to rescue ($p = 0.65$). **(C) Locomotion in the EPM in females.** Control females and 5-HT1BR KO females showed similar locomotion in the EPM across all treatment groups regardless of genotype (main interaction of genotype x drug: $F(2, 69) = 0.113$, $p = 0.89$). **(D) Home cage feeding in the NSF in males.** There was no significant effect of treatment or genotype on eating behavior in males (main interaction genotype x drug: $F(2, 55) = 0.572$, $p = 0.58$). Corticosterone had no effect on latency to eat in the home cage in both control animals (Log-Rank Mantex Cox test, chi square = 1.05, $p=0.306$) and KOs (Log-Rank Mantex Cox test, chi square = 0.226, $p=0.634$). **(E) Time spent in open arms in the EPM in males** Control males and 5-HT1BR KO males showed no effect of corticosterone treatment or drug in open arms duration in the EPM (main interaction of genotype x drug: $F(2, 62) = 1.913$, $p = 0.1562$, main effect of treatment: $F(2, 62) = 0.3593$, $p = 0.70$, main effect of genotype: $F(1, 62) = 0.33$, $p = 0.567$). **(F) Locomotion in the EPM in males.** There was no interaction of genotype by treatment in locomotion in the EPM (main interaction genotype x treatment: $F(2, 53) = 2.706$, $p = 0.08$). There was a main effect of treatment with both control and 5-HT1BR KO males showing a decrease in locomotion following corticosterone administration (main effect of treatment: $F(2, 53) = 4.309$, $p = 0.02$). KO males show decreased locomotion overall compared to control males (main effect of genotype: $F(1, 53) = 7.958$, $p = 0.01$). Error bars represent SEM, *$p < 0.05$, **$p < 0.01$.

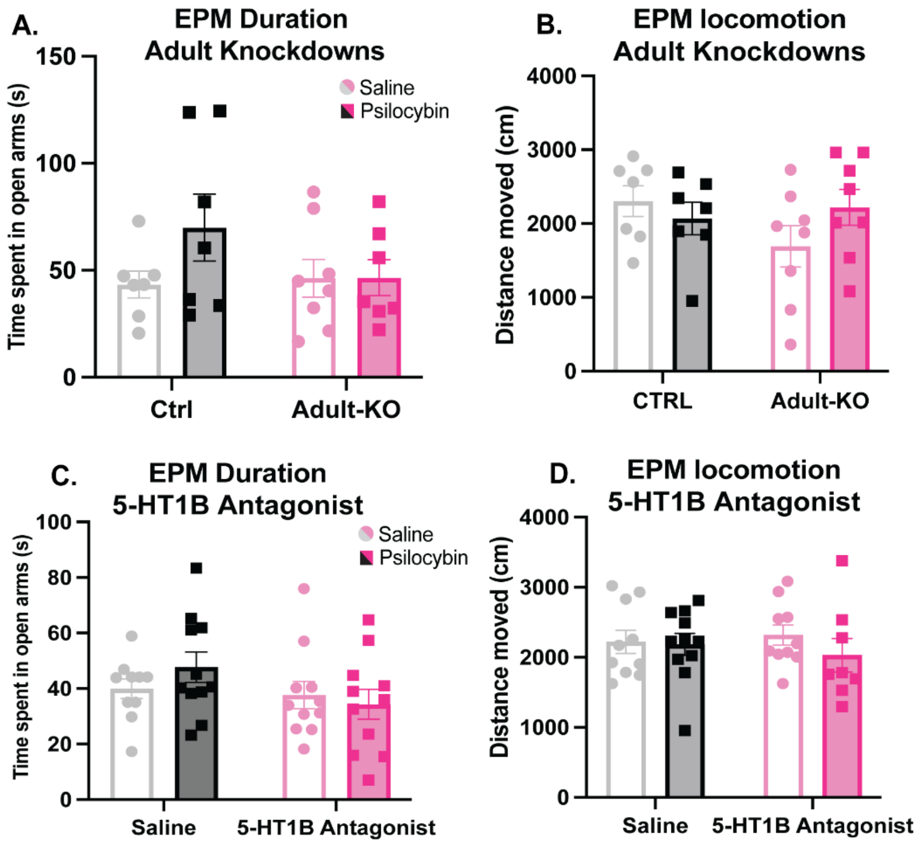


**Supplemental 5: Additional behavioral metrics in the elevated plus maze in female adult knockdowns and females treated with 5-HT1BR antagonist. (A) Duration EPM female adult knockdowns of 5-HT1BR.** Psilocybin treated control mice showed an increase in time spent in open arms when compared to psilocybin treated adult knockdowns, though not statistically significant (main interaction drug x genotype: $F(1, 25) = 1.630$, $p = 0.21$). Within controls, psilocybin increased time spent in open arms though not significantly ($t(12) = 1.582$, $p = 0.139$). Within adult knockdowns, there was no difference in entries into open arms when treated with saline versus psilocybin ($t(13) = 0.022$, $p = 0.98$). **(B) Locomotion in EPM female adult knockdowns.** All treatment groups showed similar locomotion in the EPM, regardless of treatment or genotype (main interaction of genotype x treatment: $F(1, 26) = 2.444$, $p = 0.13$). **(C) Duration EPM Female 5-HT1BR antagonist.** There was no significant effect of antagonist and drug in time spent in open arms (main interaction antagonist x drug: $F(1, 39) = 1.250$, $p = 0.27$). Psilocybin-treated controls showed a small increase in time spent in open arms in the EPM though not significant when compared to psilocybin-treated mice paired with the 5-HT1BR antagonist ($t(20) = 1.745$, $p = 0.09$). Overall, there was no effect of treatment on duration in open arms (main interaction of treatment x antagonist: $F(1, 39) = 1.250$, $p = 0.27$). **(D) Locomotion in EPM female 5-HT1BR antagonist.** All mice showed similar patterns of locomotion in the EPM regardless of treatment (main interaction of treatment x antagonist: $F(1, 35) = 0.522$, $p = 0.47$). Error bars represent SEM.

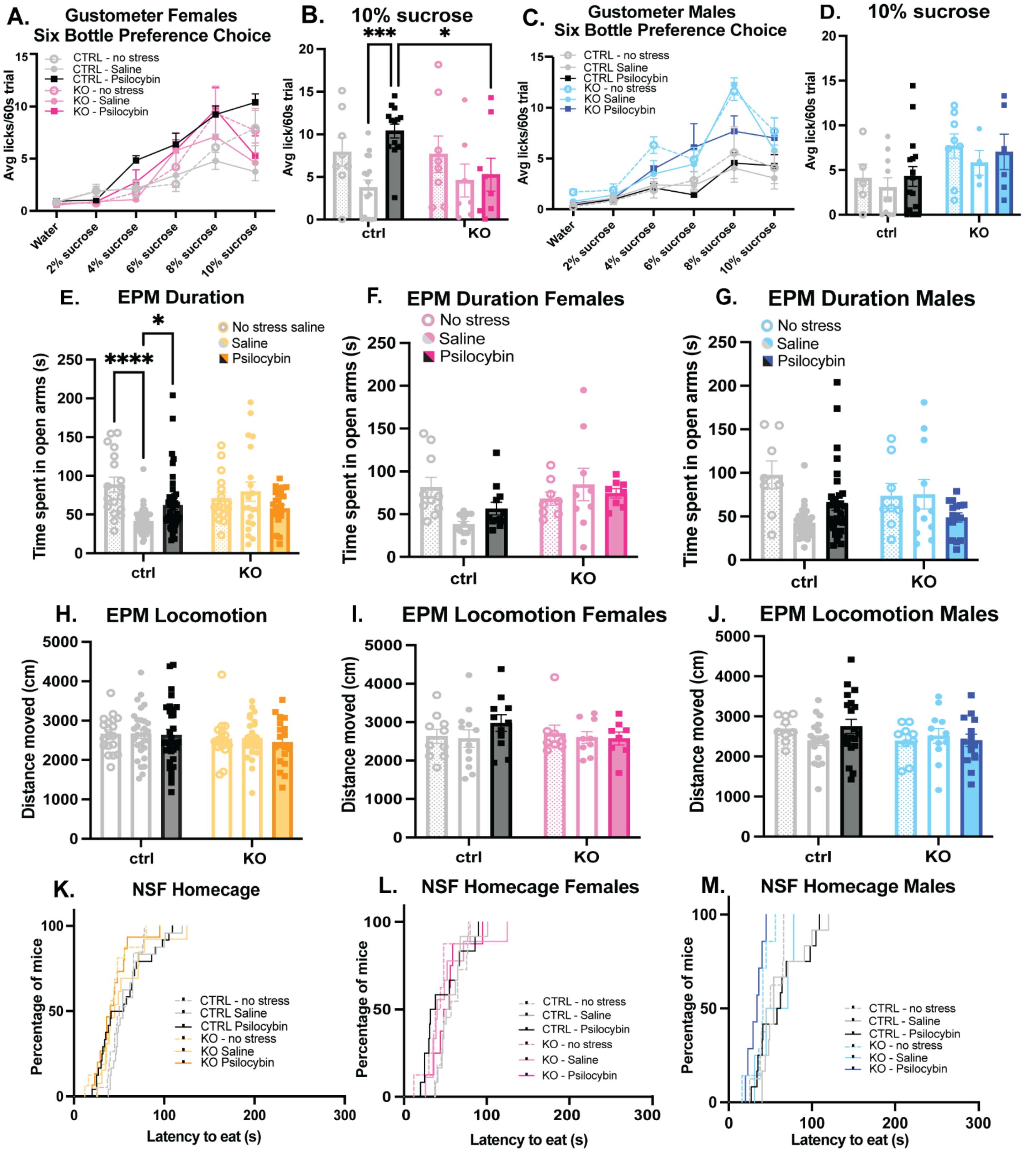


**Supplemental 6: Additional behavioral metrics in the chronic despair model (A-B) Female gustometer. (A)** There was a significant interaction of concentration, treatment, and genotype in hedonic responding (main interaction of concentration x treatment x genotype: $F(2, 276) = 3.467$, $p = 0.03$). Within control females, psilocybin rescued stress-induced anhedonia (main interaction of concentration x treatment: $F(10, 175) = 5.062$, $p < 0.0001$), but not in 5-HT1B KOs (main interaction of concentration x treatment: $F(10, 120) = 0.429$, $p = 0.93$). **(B)** There was a main effect of treatment on hedonic responding at 10% sucrose (main effect of treatment: $F(2, 55) = 4.293$, $p = 0.02$). In control females, psilocybin rescued hedonic responding at 10% sucrose ($p = 0.0004$), but not in female mice lacking 5-HT1BR ($p = 0.95$).

**(C-D) Male gustometer. (C)** Control male mice and 5-HT1BR male KOs showed no change in hedonic responding regardless of stress or drug administration (main interaction concentration x treatment x genotype: $F(2, 239) = 0.331$, $p=0.718$). Control males showed overall reduced hedonic responding relative to 5-HT1B KOs (main interaction concentration x genotype: $F(1, 239) = 15.055$). **(D)** There was no significant interaction between genotype and treatment on sucrose preference in males at 10% concentration (main interaction $F(2, 43) = 0.05$, $p=0.952$). 5-HT1BR KO males showed overall increased licking compared to control mice (main effect of genotype: $F(1, 43) = 5.68$, $p=0.022$), but there was no significant effect of treatment ($F(2, 43) = 0.46$, $p = 0.634$). **(E) Time spent in open arms in the EPM.** There was a significant interaction of genotype by treatment in time spent in open arms (main interaction of treatment x genotype: $F(2, 160) = 8.802$, $p < 0.001$). Control mice that underwent repeated forced swim spent significantly less time in open arms ($p <0\ 0.001$), which was partially rescued by psilocybin (saline vs. psilocybin: $p = 0.01$, no stress vs. psilocybin: $p = 0.02$). KO mice showed no effect of forced swim stress (no stress vs. saline: $p = 0.72$), and no effect of psilocybin either ($p = 0.10$) on time spent in open arms. **(F-G) Female and male mice showed similar patterns of duration in open arms in both control and knockout mice. (F)** There was a significant interaction of genotype by treatment in female mice (main interaction of genotype x treatment: $F(2, 53) = 4.335$, $p = 0.02$), but not of treatment ($F(2, 53) = 0.879$, $p = 0.42$). Within control females, there was a significant effect of treatment ($F(2, 31) = 7.503$, $p = 0.002$). Exposure to repeated forced swim sessions caused a significant decrease in time spent in open arms ($p = 0.002$), but this effect was attenuated in females treated with psilocybin ($p = 0.08$). KO females showed similar levels of time spent in open arms across all treatment groups ($F(2, 22) = 0.427$, $p = 0.66$), but overall spent more time in open arms than control females (main effect of genotype ($F(1, 53) = 4.220$, $p = 0.04$). **(G)** There was a significant interaction of genotype by treatment in male mice (main interaction of genotype x treatment: $F(2, 101) = 5.326$, $p = 0.01$) and a main effect of treatment ($F(2, 101) = 4.147$, $p = 0.02$), but no effect of genotype ($F(1, 101) = 0.067$, $p = 0.80$). In control males, stress decreased duration in open arms compared to non-stress males ($p = 0.0005$), which psilocybin rescued (stress-saline vs. stress-psilocybin: $p = 0.04$). In knockout males, there was no change in time spent in open arms across all treatment groups ($F(2, 31) = 1.771$, $p = 0.19$). **(H-J) Locomotion in EPM**. Control and knock out mice displayed similar locomotion in the elevated plus maze (main interaction treatment x genotype: $F(2, 134) = 0.038$, $p = 0.96$). There were no differences in locomotion across treatment or genotype (main effect of treatment: $F(2, 134) = 0.16$, $p = 0.85$; main effect of genotype: $F(1, 134) = 1.697$, $p = 0.19$). **(K-M) Home cage feeding in novelty suppressed feeding test. (K)** Overall, there was no effect of genotype or treatment on eating behavior in the home cage following NSF (Log-Rank Mantex Cox test, chi square = 2.979, $p=0.10$). **(L)** All females showed similar eating behavior in the home cage following NSF regardless of treatment or genotype (Log-Rank Mantex Cox test, chi square = 0.340, $p = 0.78$). **(M)** There was no interaction of genotype and treatment on eating behavior in the home cage following NSF in males (main interaction drug x genotype: $F(2, 44) = 1.185$, $p = 0.32$). Error bars represent SEM, $^{*}p < 0.05$, $^{**}p < 0.01$, $^{***}p < 0.001$, $^{****}p<0.0001$
